# Fibrinogen-Drug Nanoparticles Eradicate Pancreatic and Triple-Negative Breast Cancers in Mice

**DOI:** 10.64898/2026.03.18.711348

**Authors:** Rose Razavi, Michael O’Connor, James F. Hainfeld

## Abstract

Poor intratumoral drug penetration and short-lived therapeutic exposure limit durable responses to cytotoxic therapies. We developed a strategy that redirects an evolutionarily optimized clotting cascade to achieve selective, sustained drug deposition within solid tumors. A vascular disrupting agent induces tumor-specific endothelial injury and platelet activation, exposing GPIIb/IIIa. Systemically administered fibrinogen–drug nanoparticles (FDNs) bind activated platelets with multivalent avidity, amplify thrombus formation, and become immobilized within tumor vessels, creating an intratumoral drug depot that maintained paclitaxel above cytotoxic levels for over 10 days. A single 15-minute treatment eradicated advanced triple-negative breast tumors in 88% of mice, while docetaxel-based FDNs produced durable eradication of pancreatic tumors in all treated mice (9/9). These findings establish a clinically translatable clot-guided platform for sustained therapeutic exposure in refractory cancers.

## Introduction

Despite major advances in oncology, many solid tumors remain resistant to durable control. Classical cytotoxic agents suffer from systemic toxicity, rapid elimination and drug resistance(*1, 2*), while nanoparticle formulations—though designed to improve solubility, circulation time, and biodistribution—still deliver only a small fraction of drug to tumors. Across dozens of studies, nanoparticles achieve a median of 2.2% of the injected dose per gram of tumor(*3*), reflecting poor extravasation, limited stromal penetration, and rapid clearance (*4*). Even when nanoparticles reach tumors, ineffective release kinetics and poor cellular uptake further limit efficacy (*5,6*). Immunotherapies and antibody–drug conjugates provide greater specificity(*7*), yet many tumors remain immunologically cold(*8, 9*), rapidly develop resistance, or lack the required surface markers(*10*). Collectively, these limitations underscore the persistent challenge of achieving high, sustained therapeutic exposure within solid tumors.

Several groups have previously explored strategies that leverage coagulation biology or therapy-induced vascular changes to enhance nanoparticle accumulation in tumors. Prior approaches have used coagulation activation or vascular disruption to enhance nanoparticle accumulation (*12*-*15*). These studies demonstrated that therapeutic modulation of the tumor microenvironment can enhance nanoparticle localization.

However, these prior systems primarily relied on ligand-mediated targeting or externally induced signaling mechanisms and did not directly harness fibrinogen—the endogenous substrate of platelet aggregation—to amplify thrombosis and create a sustained intratumoral drug reservoir. In contrast, the strategy described here uses fibrinogen itself as both the nanoparticle scaffold and the targeting element, enabling direct multivalent binding to activated platelet GPIIb/IIIa receptors and incorporation into the growing platelet–fibrin network. This design allows the drug-loaded particles to become physically integrated into tumor clots, producing both vascular embolization and prolonged intratumoral drug release.

To overcome these anatomical and physiological barriers, we turned to an evolutionarily refined biological system already specialized for rapid, localized polymerization: the clotting cascade(*16*). When a blood vessel is injured, platelets rapidly activate, expose the high-affinity integrin GPIIb/IIIa(*17*), bind fibrinogen **(Fig. 1B)**, and assemble a multivalent platelet-fibrin matrix that seals vascular breaches with remarkable speed and strength. This process relies on high-avidity binding, explosive amplification, and site-specific polymer formation, precisely the attributes required for effective tumor-localized drug delivery.

**Fig. 1.**
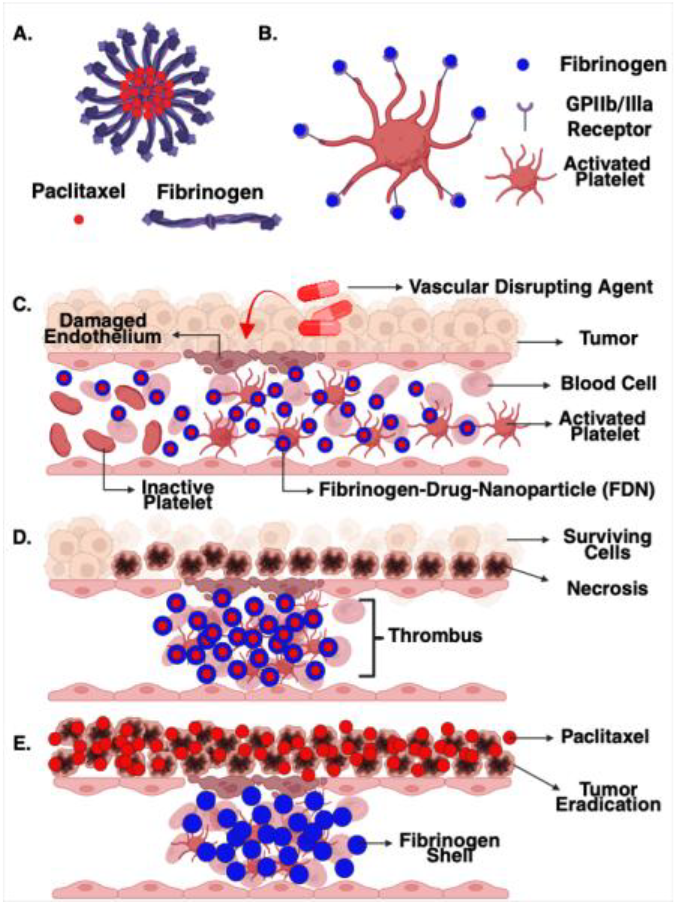
Mechanism of fibrinogen-drug nanoparticle. **(FDN)** therapy **(A)** Schematic structure of an FDN, composed of paclitaxel encapsulated within a fibrinogen-based nanoparticle. **(B)** Activated platelets expose the integrin GPIIb/IIIa, which binds the fibrinogen component of FDNs with high avidity. **(C)** A vascular disrupting agent selectively injures tumor endothelium, triggering local platelet activation and enabling intravascular capture of circulating FDNs. **(D)** Multivalent FDN binding amplifies platelet aggregation and fibrin polymerization, forming an obstructive thrombus that embolizes tumor vessels and induces downstream necrosis. **(E)** Immobilized FDNs act as a drug-releasing depot, delivering paclitaxel over extended periods to eradicate surviving tumor cells.

We hypothesized that if a tumor could be primed to transiently mimic vascular injury, activated platelets could be exploited as a drug-capturing surface. This approach integrates three coordinated steps **(Fig. 1C-E)**: (1) targeted endothelial damage to activate platelets, (2) fibrinogen–drug nanoparticle (FDN) binding that amplifies thrombosis and embolizes tumor vessels, and (3) sustained intratumoral drug release from the resulting platelet–fibrin scaffold. Together, these steps convert the tumor’s own vasculature into a localized delivery and destruction platform.

Mechanistically, FDNs out-compete endogenous fibrinogen for GPIIb/IIIa binding; their multivalency and larger size drive more extensive platelet crosslinking and more robust vessel occlusion, inducing downstream necrosis and enabling prolonged intratumoral drug exposure. This approach achieves sustained intratumoral drug exposure and durable tumor eradication.

## Results

### Construction and Characterization of Fibrinogen–Drug Nanoparticles

Sonicating fibrinogen with paclitaxel produced uniform 183 ± 8.9 nm nanoparticles with low polydispersity (PDI, 0.18 ± 0.04, **Fig. 2A**) and high encapsulation efficiency (91.7 ± 4.0%). Given their larger size and multimeric structure (**Fig. 1A**), FDNs were expected to undergo more extensive fibrin polymerization than free fibrinogen once exposed to thrombin. To evaluate this, we used the Clauss assay, in which thrombin converts fibrinogen to fibrin(*18*) and the resulting increase in turbidity reflects polymer formation over time(*19*). Consistent with this prediction, FDNs generated ∼30-fold greater fibrin polymerization than an equivalent amount of free fibrinogen (**Fig. 2B**), demonstrating markedly enhanced incorporation into clot formation and supporting their ability to amplify thrombosis in vivo.

**Fig. 2.**
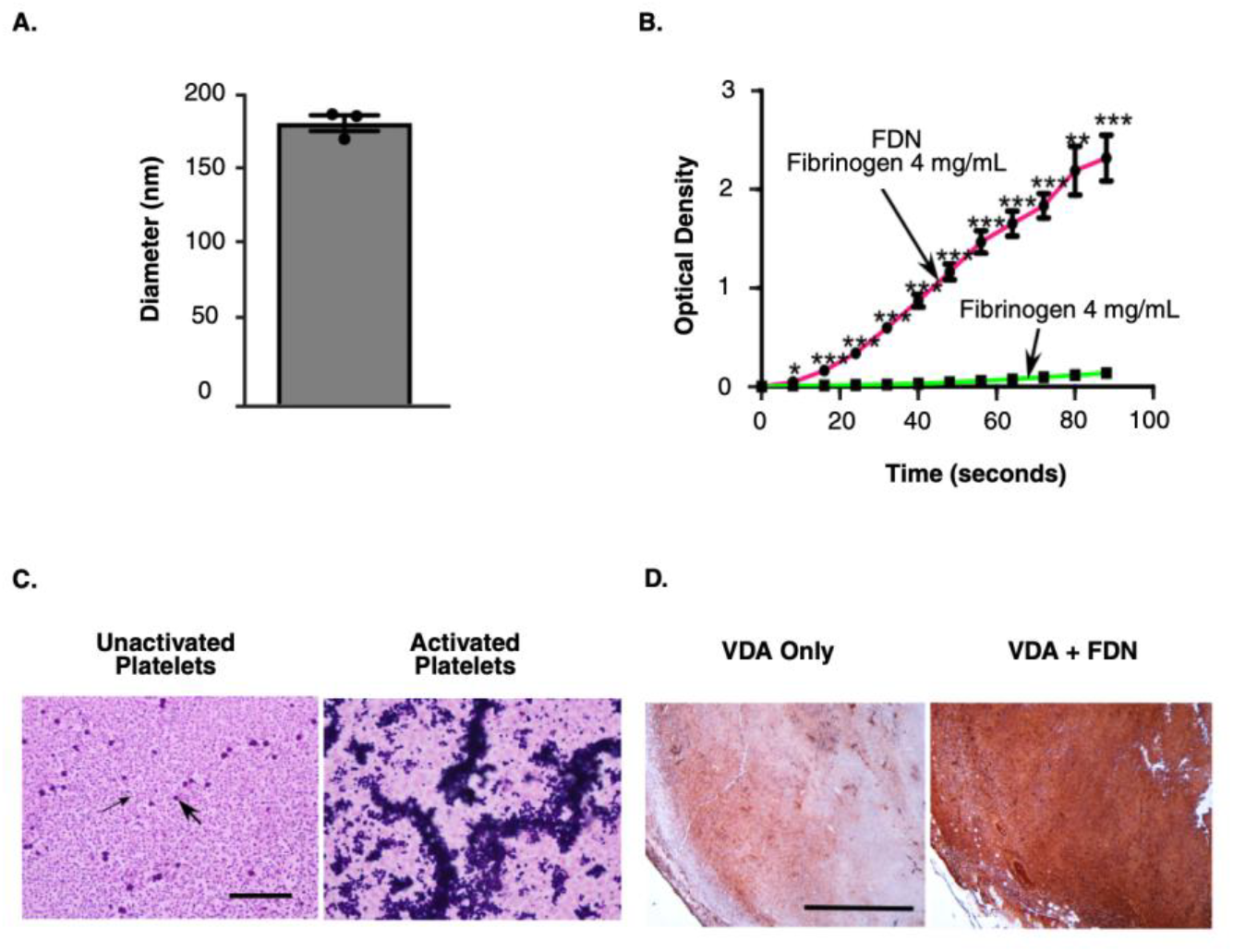
Fibrinogen–drug nanoparticles retain coagulation activity, promote platelet aggregation, and enhance intratumoral fibrinogen deposition following vascular disruption. **(A)** Hydrodynamic diameter of fibrinogen–drug nanoparticles (FDNs), demonstrating uniform nanoparticle size. Bars represent mean ± SEM. **(B)** Clauss fibrin polymerization assay comparing free fibrinogen and FDNs containing an equivalent fibrinogen concentration (4 mg/mL) following thrombin activation. FDNs exhibit markedly enhanced polymerization kinetics and optical density relative to free fibrinogen, indicating preserved and amplified coagulation activity (*p* < 0.0001). **(C)** Platelet aggregation in platelet-rich plasma following exposure to FDNs. Left, platelets isolated in the presence of prostaglandin E1 (aggregation inhibited) show no aggregation upon FDN addition. Right, ADP-activated platelets rapidly form large aggregates upon FDN exposure. Large arrow indicates red blood cell; small arrow indicates platelet. Scale bar, 20 μm. **(D)** Immunohistochemical staining for fibrinogen (brown) in EMT6 tumors 24 h after treatment with vascular disrupting agent (VDA) alone or VDA combined with FDNs. VDA + FDN treatment results in extensive fibrinogen accumulation throughout tumor tissue compared with VDA alone, indicating enhanced nanoparticle deposition at sites of vascular damage. Scale bar, 0.5 mm.

### FDNs Bind Selectively to Activated Platelets

FDNs were next evaluated for their ability to bind specifically to activated platelets. To do this, platelet-rich plasma (PRP) was isolated from mouse blood and used in two parallel assays. In one condition, prostaglandin E1 was added to PRP to inhibit platelet activation; when mixed with FDNs, no binding or aggregation was observed (**Fig. 2C**), indicating minimal interaction with quiescent platelets. In contrast, when PRP was activated with ADP before the addition of FDNs, extensive and immediate platelet aggregation occurred (**Fig. 2C**). These findings demonstrate that FDNs selectively interact with activated platelets—consistent with GPIIb/IIIa-mediated recognition—and support their ability to target platelet activation sites generated after tumor endothelial injury.

### VDA-Induced Platelet Activation Enables Intratumoral Capture of Fibrinogen Constructs

Administration of the vascular disrupting agent DMXAA selectively damaged the tumor vasculature, leading to rapid platelet activation. Immunohistochemistry revealed substantially greater fibrinogen deposition throughout tumors treated with VDA + FDNs than with VDA alone, demonstrating efficient intratumoral capture of the nanoparticles. When tumors were treated with VDA alone, fibrinogen deposits were typically restricted to the tumor periphery, where VDAs preferentially disrupt angiogenic endothelium (*20*– *22*). Surprisingly, when FDNs were used in combination with VDA, heavy fibrinogen deposits were observed not only at the periphery but also deeper within the tumor mass (**Fig. 2D**). This widespread intratumoral deposition is consistent with multivalent FDN– platelet interactions amplifying thrombus propagation, allowing clot formation to extend into deeper tumor regions.

### Fibrinogen–Paclitaxel Nanoparticles (FDN-PTX) Drive Durable Responses in a Triple-Negative Breast Cancer Model

Within 24–48 hours of VDA + FDN treatment, tumors became visibly black and necrotic, far exceeding the mild peripheral necrosis observed with VDA alone (**Fig. 3A**). This therapy was accompanied by transient weight loss over the first five days, after which animals rapidly recovered to baseline (**Fig. 3G**). These highly necrotic tumors regressed over subsequent weeks by 4–5 weeks after treatment, leaving no detectable residual tumor (**Fig. 3B,C**). This vascular shutdown produced robust therapeutic efficacy across a wide range of tumor sizes (145–1020 mm^3^; mean 543 mm^3^, **Fig. 3F**). A single treatment resulted in complete tumor regression in 14/16 mice with no recurrence over 200 days (**Fig. 3D,E**).

**Fig. 3.**
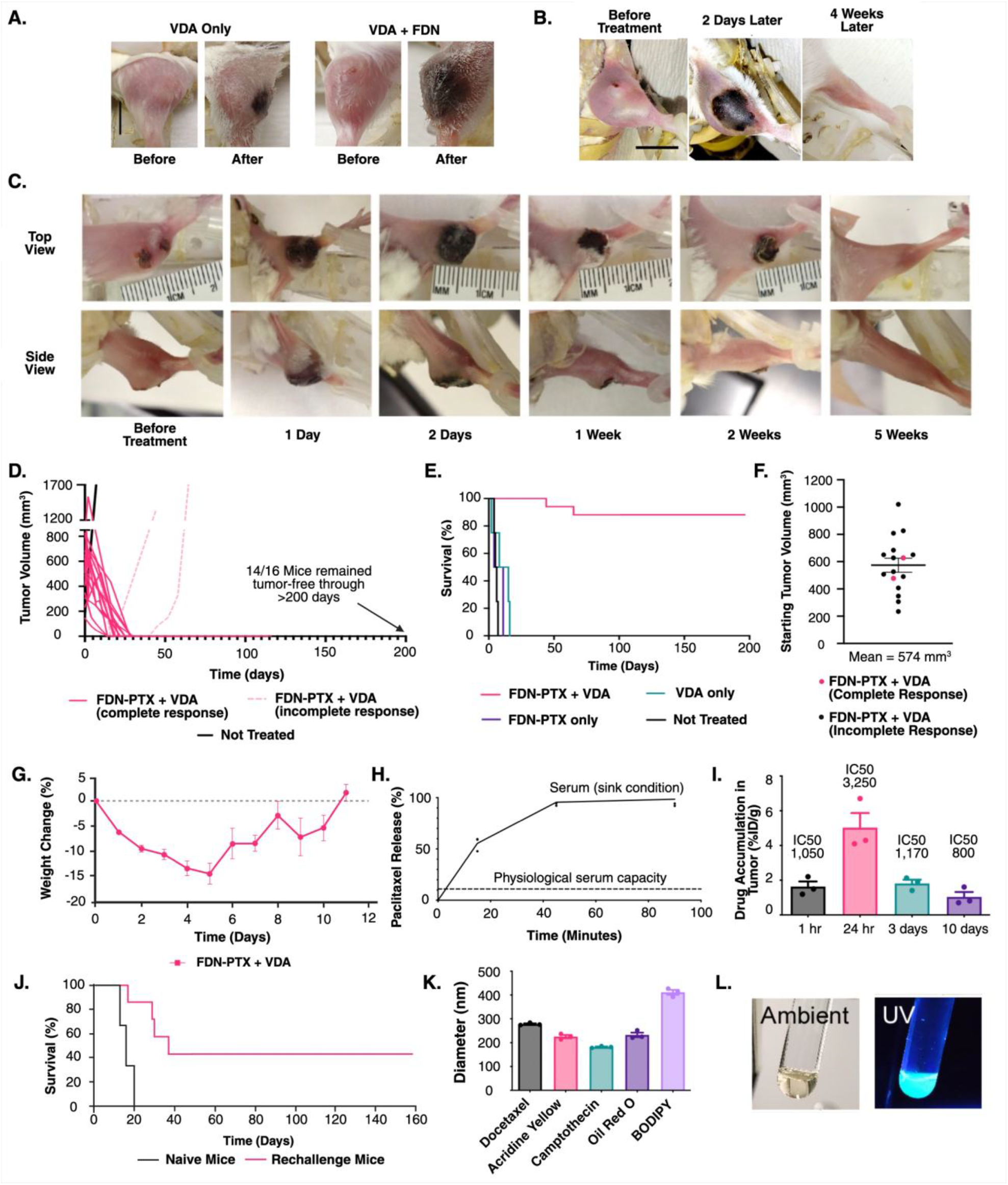
Fibrinogen–drug nanoparticles (FDNs) combined with vascular disrupting agents induce rapid tumor necrosis, durable cures, and long-term antitumor immunity in EMT6 TNBC. **(A)** Representative images of EMT6 triple-negative breast tumors before and 24 h after treatment with vascular disrupting agent (VDA) alone or VDA combined with fibrinogen–drug nanoparticles (FDNs). Combined VDA + FDN treatment induces markedly greater tumor necrosis compared with VDA alone. Scale bar, 0.5 cm. **(B)** Longitudinal photographs of a large 1020 mm^3^ EMT6 tumor before treatment, 2 days after a single VDA + FDN treatment, and 4 weeks later, showing complete tumor elimination without regrowth. Scale bar, 1 cm. **(C)** Serial top- and side-view images illustrating treatment response over time following VDA + FDN therapy. Extensive tumor necrosis and scab formation occur within days, followed by progressive tumor shrinkage and complete resolution by 5 weeks, with restoration of normal tissue appearance and no detectable recurrence. **(D)** Individual tumor volume trajectories following VDA + FDN therapy. Each line represents one mouse. Fourteen of sixteen mice exhibited complete and durable tumor regression, remaining tumor-free for >200 days. Typical tumor growth of an untreated control is shown for comparison. **(E)** Kaplan–Meier survival analysis of EMT6 tumor–bearing mice. VDA + FDN treatment resulted in 88% long-term survival (>200 days), whereas all control groups (untreated, FDN only, or VDA only) succumbed within 20 days. Survival of the VDA + FDN group was significantly improved compared with all controls (log-rank Mantel–Cox test, *p* < 0.001). Group sizes: VDA + FDN, *n* = 16; untreated, *n* = 4; VDA only, *n* = 4; FDN only, *n* = 4. **(F)** Distribution of starting EMT6 tumor volumes at the time of treatment, demonstrating effective tumor eradication across a broad range of initial tumor sizes. **(G)** Body weight changes following VDA + FDN therapy, showing transient weight loss with recovery to baseline within ∼11 days, indicating tolerable systemic toxicity. **(H)** Paclitaxel (PTX) release from FDNs into serum under sink conditions. PTX is released nearly to completion within 1 h. The dashed line indicates the estimated low release into the blood pool after injection due to poor serum capacity to solubilize a hydrophobic drug. **(I)** Tumor accumulation of PTX following VDA + FDN administration, measured at 1 h, 24 h, 3 days, and 10 days post-injection, demonstrating rapid tumor delivery and sustained intratumoral drug levels. **(J)** Re-challenge experiment demonstrating treatment-induced antitumor immunity. Mice cured by VDA + FDN therapy either rejected secondary tumor implantation or exhibited significantly delayed tumor growth compared with naïve controls (log-rank Mantel–Cox test, *p* = 0.0048). **(K)** Hydrodynamic diameters of fibrinogen nanoparticles loaded with diverse small-molecule cargos, demonstrating modular nanoparticle construction. **(L)** Representative images of fluorescently labeled fibrinogen nanoparticles under ambient light and UV illumination.

### Sustained High Intratumoral Paclitaxel Levels Following FDN Delivery

FDNs delivered ∼5 %ID/g paclitaxel at 24 h, exceeding the IC50 by ∼3,000-fold and remaining above cytotoxic levels for >10 days, yielding an AUC of 5,620 h·µg/g (Fig. 3I). When initially injected, only ∼11% of the paclitaxel can be released into the serum due to its poor solubility in serum (*24*) (**Fig. 3H**). Under sink conditions, paclitaxel release from fibrin-clotted FDNs was rapid reaching completion within ∼1 hour (**Fig. 3H**). This concentration-ratio dependent release, combined with the physical immobilization of FDNs within tumor thrombi, provides a mechanistic explanation for the sustained high drug levels in tumors.

### FDN Therapy Induces Tumor-Specific Immunity

Immunotherapy is a highly promising strategy for long-term tumor control; however, many tumors exhibit an immunologically “cold” phenotype and fail to respond (*8, 9*). To determine whether FDN therapy stimulates a protective antitumor immune response, mice that had been treated and cured were rechallenged with tumor cells in the contralateral leg, and tumor growth was monitored (**Fig. 3J**). Notably, 43% (3 of 7) of previously treated mice showed complete absence of tumor growth, while an additional 4 of 7 exhibited substantially slower tumor progression compared to naïve controls, indicating that FDN therapy induces an adaptive immune response capable of suppressing tumor recurrence.

### Platform Versatility for Multiple Hydrophobic Agents

Since paclitaxel readily incorporates into fibrinogen, we next evaluated five additional hydrophobic compounds to assess platform versatility. Multiple hydrophobic compounds, including chemotherapeutics, dyes, and fluorophores, formed fibrinogen-based nanoparticles comparable to FDN-PTX (**Fig. 3K**), demonstrating platform versatility. This versatility suggests broad translational potential, enabling adaptation of the platform for both therapeutic and diagnostic applications across multiple disease contexts.

### Fibrinogen-Docetaxel Nanoparticles (FDN-DTX) Drive Durable Responses in a Desmoplastic Pancreatic Cancer Model

Having established robust efficacy in the EMT6 breast cancer model and demonstrated compatibility with multiple hydrophobic payloads, we next evaluated the platform in a KPC pancreatic cancer model, a clinically relevant and therapeutically refractory tumor context characterized by dense desmoplasia and dysregulated coagulation biology. The KPC model closely recapitulates key genetic drivers and pathological features of human pancreatic ductal adenocarcinoma (PDAC), including activating Kras and mutant p53 signaling (*30*–*32*).

Initial evaluation of paclitaxel-loaded fibrinogen drug nanoparticles (FDN-PTX) in KPC tumors demonstrated measurable antitumor activity but produced heterogeneous outcomes. Tumor growth curves revealed substantial variability in therapeutic response, with 2 of 13 treated animals achieving complete tumor regression while others exhibited partial or transient responses followed by tumor progression (**Supplementary Fig. 1A**). Consistent with these observations, Kaplan–Meier survival analysis showed improved survival relative to untreated controls but incomplete long-term tumor control (**Supplementary Fig. 1B**).

To determine whether an alternative taxane payload could improve therapeutic performance in this tumor context, we evaluated docetaxel (DTX), a related taxane with higher affinity for tubulin (*33*), using a modified formulation procedure that yielded a larger fibrinogen drug nanoparticle with reduced colloidal stability incorporating DTX (FDN-DTX) (433 ± 137 nm with a PDI of 0.21 ± 0.06) immediately after preparation) that increased in size to 575 ± 124 nm with a PDI of 0.29 ± 0.05 after 20 minutes.

With this optimized preparation, VDA-primed delivery of FDN-DTX produced rapid tumor regression and durable complete responses in all treated animals (9/9). Tumors became undetectable by caliper measurement within approximately 12–25 days following treatment and remained absent throughout the duration of the study. Mean tumor growth curves (± SEM) demonstrate complete and sustained tumor clearance in the FDN-DTX + VDA group, whereas untreated and VDA-only control groups exhibited progressive tumor growth with overlapping trajectories (**Fig. 4A**). Kaplan–Meier analysis further demonstrated durable survival benefits, with all treated animals remaining tumor-free beyond 100 days of observation (**Fig. 4B**). Treatment caused transient ∼10–12% weight loss followed by recovery (**Fig. 4C**). Therapeutic responses occurred across a broad range of tumor volumes (100–>1000 mm^3^; mean 454 mm^3^; **Fig. 4D**). Together, these findings demonstrate that docetaxel-based fibrinogen nanoparticles markedly enhance therapeutic efficacy in KPC pancreatic tumors.

**Fig. 4.**
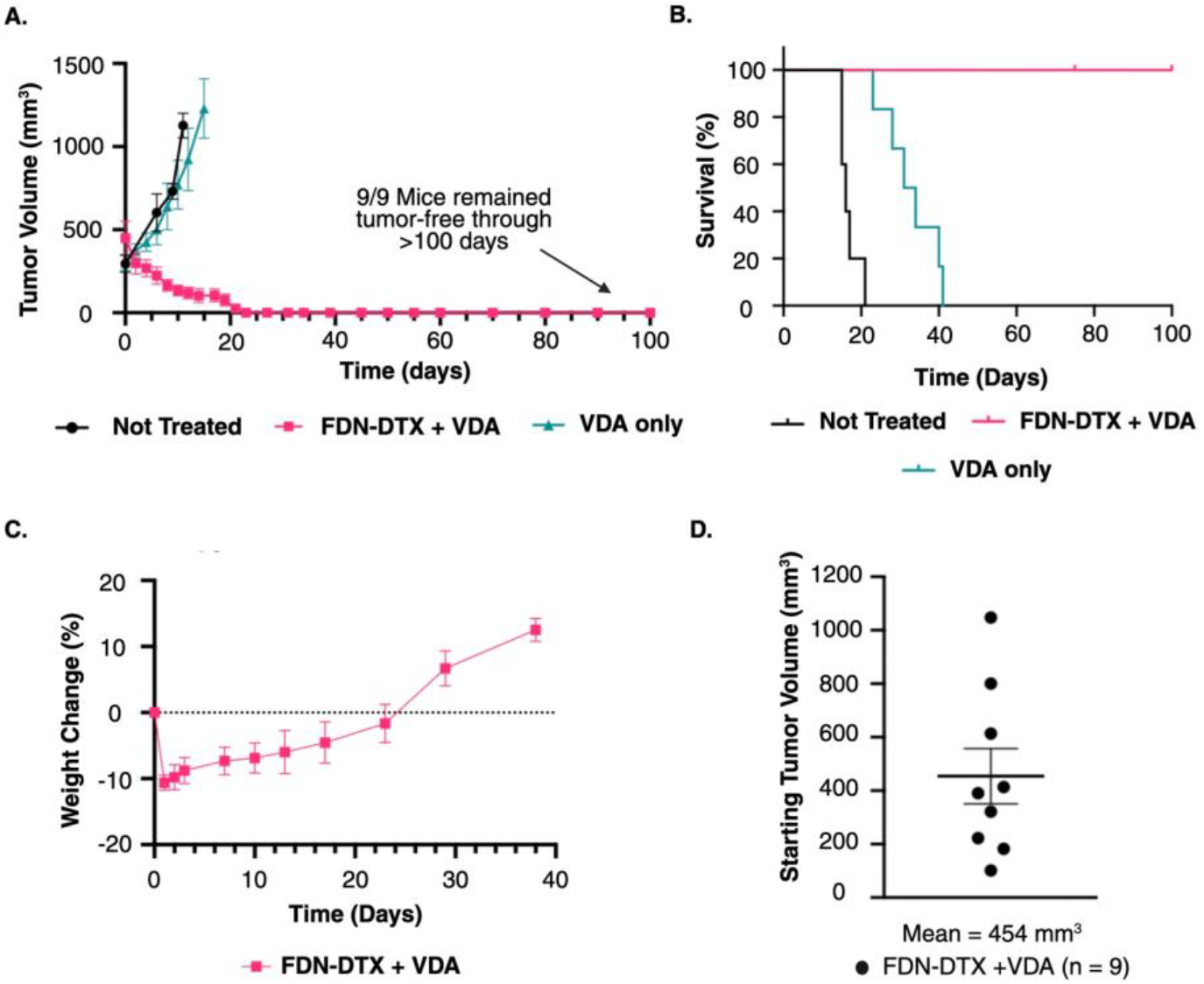
Fibrinogen-docetaxel nanoparticles eradicate KPC pancreatic tumors following vascular priming. **(A)** Tumor volume over time in KPC pancreatic tumor– bearing mice treated with the indicated conditions: untreated (n = 5), vascular disrupting agent (VDA) alone, or VDA + FDN-DTX (n = 9). Combination therapy produced rapid tumor regression and complete disappearance of tumors within ∼12–25 days, whereas untreated and VDA-only groups showed progressive tumor growth. Data represent mean ± SEM. **(B)** Kaplan–Meier survival analysis of treated mice. VDA + FDN-DTX produced durable survival, with all treated animals remaining tumor-free throughout the observation period (>100 days), whereas untreated controls succumbed rapidly to tumor progression. **(C)** Body weight changes following treatment. VDA + FDN-DTX caused a transient decrease in body weight (∼10–12%) followed by recovery to baseline levels. **(D)** Distribution of starting tumor volumes at treatment initiation in the VDA + FDN-DTX cohort (n = 9). Tumors ranged from ∼100 to >1000 mm^3^ (mean = 454 mm^3^), demonstrating responses across a wide range of initial tumor sizes.

## Discussion

This study presents a fundamentally different strategy for cancer drug delivery: hijacking an evolutionarily optimized hemostatic cascade to concentrate cytotoxic agents directly within tumor vasculature. Unlike conventional nanoparticle systems (typically optimized to be 50–200 nm to facilitate extravasation and tumor penetration (*43, 44*)), FDNs exploit the exposed activated platelets within tumor blood vessels, eliminating the need for interstitial transport and allowing the use of larger, aggregation-capable constructs. Larger particle–platelet complexes increase embolic potential, enhancing vessel occlusion.

Fibrinogen–platelet interactions drive this mechanism with exceptional affinity. The fibrinogen–GPIIb/IIIa interaction (Kd ≈ 15 nM), comparable to high-affinity antibodies (*45, 46*), becomes effectively irreversible through multivalent binding (*45, 47*). This interaction initiates platelet activation and aggregation (*48*), amplifying thrombus formation. Resulting ischemia leads to apoptosis and necrosis (*49*), providing an advantage over conventional therapies that rely primarily on apoptosis (*50*); necrosis is particularly effective against tumor cells that have evolved apoptosis resistance.

This embolic trapping creates a stable, long-acting drug reservoir. FDNs delivered 5 %ID/g paclitaxel at 24 hours, substantially higher than many reported delivery systems, including Abraxane (∼0.1 %ID/g (32,33) and liposomes (∼0.06 %ID/g (34)). Although modern antibody–drug conjugates can reach ∼45 %ID/g (*7*), the critical distinction lies in sustained local exposure. With FDNs, paclitaxel levels remained above the IC50 for >10 days, resulting in an AUC of 5,620 h·µg/g, nearly 100-fold greater than values typically reported for antibody-drug conjugates (*51*). Importantly, prolonging paclitaxel exposure from 24 to 72 hours has been shown to increase cytotoxicity by up to 200-fold (*52*). Such prolonged retention is facilitated by nanoparticle entrapment within platelet–fibrin matrices, limiting systemic clearance and enabling slow release precisely within the tumor microenvironment. Paclitaxel’s limited serum solubility restricts systemic release, while intratumoral retention (>10 days; **Fig. 3I**) and fibrin-mediated entrapment enable sustained local drug exposure.

The durable eradication of aggressive KPC pancreatic tumors further highlights the strength and generalizability of this approach. Even large established tumors exceeding 1000 mm^3^ were eliminated following VDA-primed FDN–DTX treatment, demonstrating that this strategy can overcome the highly desmoplastic and therapy-resistant microenvironment characteristic of pancreatic ductal adenocarcinoma. The rapid and complete responses observed across treated animals suggest that coupling vascular disruption with fibrinogen-mediated drug deposition enables highly efficient intratumoral drug retention and associated cytotoxicity, even in tumor types traditionally resistant to chemotherapy. These findings support the potential of this platform, particularly in combination with vascular priming strategies, for application across multiple solid tumor types.

A potential explanation for differences in response to vascular priming between EMT6 and KPC tumors may relate to underlying variations in tumor-associated coagulation and platelet biology. Pancreatic ductal adenocarcinoma (PDAC) exhibits elevated thrombosis and platelet activation compared to breast cancer, which may enhance fibrinogen-based nanoparticle capture following vascular disruption (*34*–*42*).

Mechanistic studies further demonstrate that tumor cells can induce platelet activation and aggregation through tissue factor– and thrombin-dependent pathways (*39*–*41*), and intratumoral platelets are increasingly recognized as active components of the tumor microenvironment (*42*). Together, these observations support the plausibility that pancreatic tumors may exhibit a heightened baseline propensity for platelet activation and fibrin/platelet capture, which could enhance localization and activity of fibrinogen-based constructs following vascular disruption.

This vascular-targeted approach offers multiple advantages. Because it does not rely on tumor-specific biomarkers, it has broad applicability across cancer types. In addition, targeting endothelial cells helps circumvent multidrug resistance, as these cells are less prone to acquiring mutations than tumor cells (*55*). The system is composed of biologically compatible components, namely fibrinogen and platelets, which may support tolerability.

Although DMXAA reached phase III clinical testing, it ultimately failed to improve overall survival (*56*), likely due to two key limitations. First, DMXAA functions as a STING agonist in mice but not in humans (*56*). Second, although DMXAA reduced tumor blood flow by up to 66% (*57*), residual tumor cells at the periphery survived by obtaining oxygen and nutrients from adjacent normal vasculature, an effect known as the “viable rim” phenomenon (*58*). In contrast, the clot-based FDN strategy may overcome these limitations by coupling vascular disruption with prolonged intratumoral drug release, ensuring cytotoxic exposure extends to peripheral tumor regions. Thus, while DMXAA itself is not clinically viable, the underlying mechanistic concept may still be leveraged using synthetic STING agonists or alternative vascular-disrupting agents in combination with the FDN platform (*59, 60*). Vascular disruption can also be achieved by radiation (*42*), heat (*61*), mechanical stress (*62*) (e.g., lithotripsy), or other means.

Importantly, EMT6 tumors are considered immunologically “cold” and poorly responsive to immune activation (*63*). However, our rechallenge data indicate that FDN therapy may induce immunological memory: 43% (3/7) of cured mice exhibited complete tumor rejection upon contralateral rechallenge, and 4/7 showed markedly slower tumor growth, suggesting the generation of a sustained adaptive response. Necrotic tumor cell death likely contributes via release of inflammatory cytokines and immunostimulatory factors (*50*), potentially enhancing antitumor immunity and facilitating abscopal-like effects (*64, 65*).

Personalization requirements often limit access to advanced cancer therapies; however, more than 90% of all cancers rely on neovascularization (*66*), making this strategy broadly relevant. Additional hydrophobic agents—including camptothecin, docetaxel, oil red O, acridine yellow and BODIPY—successfully formed nanoparticles like FDN–paclitaxel (**Fig. 3K**), demonstrating formulation extensibility.

While highly promising, this approach carries potential safety concerns that warrant further investigation. Because the clotting system is co-opted for drug targeting, there is a risk of off-target thrombotic events, which could lead to adverse vascular outcomes such as stroke or myocardial infarction. Fibrin deposition is also implicated in several pathological conditions (*67*), and patients experiencing extensive tissue damage, such as burn victims, may exhibit hypercoagulability (*68*), highlighting the importance of careful toxicity evaluation. Additionally, although the vascular-disrupting agent DMXAA is effective in mice, it lacks full activity in humans; however, synthetic STING agonists and alternative vascular-disrupting agents are currently under exploration (*59, 60*).

Chemotherapy fails in a large fraction of cases, often due to the development of tumor resistance to anticancer agents (*69*). Because tumor eradication was achieved after a single treatment over a relatively short time (∼3-5 weeks), the opportunity for tumors to develop resistance may be reduced.

The platform may be extended using alternative hydrophobic drugs or platelet-targeting ligands and integrated with existing nanoparticle systems or vascular-priming strategies.

A surprising result was that only a single, simple, non-invasive ∼15-minute treatment was necessary to fully eradicate tumors compared to the extensive, multiple, prolonged and often less effective treatments currently available. We attribute this to the multifunctionality of fibrinogen–drug nanoparticles, which combine avid targeting, polymerization and stabilization within tumors, vessel embolization, sustained drug release, and biodegradability.

Together, these results establish a clinically translatable clot-guided drug delivery platform that converts tumor vasculature into a long-acting intratumoral drug depot. By co-opting evolution’s fastest polymerizing system, this strategy integrates vascular embolization, sustained chemotherapy delivery, and immune activation into a unified therapeutic approach for drug-resistant solid tumors.

## Conclusion

We present a clot-guided therapeutic strategy that harnesses the clotting cascade to eradicate tumors. By combining tumor-selective vascular disruption with fibrinogen–drug nanoparticles, this strategy triggers a self-amplifying intratumoral thrombosis that both blocks tumor blood flow and creates a long-acting drug reservoir. These coordinated steps—(1) targeted endothelial injury, (2) nanoparticle-driven platelet capture and vessel embolization, and (3) sustained intratumoral cytotoxic release—convert the tumor vasculature into a therapeutic delivery and destruction platform.

Using this approach, fibrinogen–taxane nanoparticles produced durable eradication of aggressive solid tumors in two preclinical models. In triple-negative breast cancer (EMT6), a single, noninvasive treatment combining DMXAA with fibrinogen–paclitaxel nanoparticles induced extensive tumor necrosis and sustained drug exposure for more than 10 days, resulting in complete tumor elimination in most animals. Extending the platform to a highly desmoplastic KPC pancreatic cancer model using fibrinogen–docetaxel nanoparticles produced rapid and durable tumor clearance even in large established tumors.

Together, these findings establish a first-in-class clot-guided drug delivery strategy that converts tumor vasculature into a long-acting intratumoral drug depot. This approach integrates vascular emobilization, sustained drug delivery, and immune activation into a broadly applicable therapeutic platform.

## Acknowledgments

We would like to thank Yaroslav Stanishevskiy for help with cell culture, Keri Fitzgerald for mouse husbandry and maintenance, and Dolly Shaw for mouse care and help with tumor measurements. All figures were created on Biorender.com.

## Author contributions

Conceptualization: RR, JFH

Methodology: RR, MO, JFH

Investigation: RR, MO, JFH

Visualization: RR, MO, JFH

Funding acquisition: JFH

Project administration: JFH

Supervision: JFH

Writing – original draft: JFH

Writing – review & editing: RR

## Data, code, and materials availability

All data are available in the main text or the supplementary materials.

