## supplemental information for "Fibrinogen-Drug Nanoparticles Eradicate Pancreatic and Triple-Negative Breast Cancers in Mice"

**The PDF file includes:**

Materials and Methods
Figs. S1
References

### Materials and Methods

Materials

Fibrinogen (Fib 1, plasminogen-depleted; bovine and human) was obtained from Enzyme Research Laboratories (South Bend, IN). Docetaxel (DTX) and paclitaxel (PTX) were purchased from LC Laboratories (Woburn, MA). DMXAA was obtained from Tokyo Chemical Industry Co. (D5235) or InvivoGen (tlrl-dmx) and prepared as a 2 mg/mL solution in 0.1 M Tris buffer (pH 8.0). Polyethylene glycol (PEG 200, MW) was purchased from Thermo Fisher Scientific (B21918). Poly-L-lysine (PB920), ADP, bovine serum albumin (BSA; A7906), ethyl acetate (34858), and butyl 4-hydroxybenzoate (54680) were obtained from Sigma-Aldrich. Wright–Giemsa stain was purchased from Volu-Sol, and Permount mounting medium was obtained from Fisher Scientific. Thrombin was purchased from MP Biomedicals (154163), and prostaglandin E1 was obtained from Tokyo Chemical Industry. An ultrasonic disruptor (Misonix SL-2000) equipped with a microtip probe was used for nanoparticle preparation. Dynamic light scattering (DLS) measurements were performed using a Brookhaven Instruments 90Plus particle size analyzer. BALB/c mice were obtained from Taconic Biosciences (Rensselaer, NY). C57/BL (B6(Cg)-Tyr<c-2J>/J ) mice were from The Jackson Laboratory (Bar Harbor, ME). EMT6 murine mammary carcinoma cells were obtained from the American Type Culture Collection (ATCC). The KPC cell line was kindly provided by Dr. Mario Shields, Pathology Department, Stony Brook University, Stony Brook, NY) which was generated from the KPC (KrasG12D/+; Trp53R172H/+; P48-Cre) mouse model (Jackson Laboratory #014647).

Methods

***Fibrinogen-Paclitaxel Nanoparticle (FDN-PTX) Preparation:*** FDN-PTX was prepared using a sonication-based formulation method. In a typical synthesis, 26.7 mg fibrinogen was dissolved by adding 125 µL of 0.1M borate buffer pH 8.6, then 50 µL 3M NaCl, then 660 µL water without mixing and incubated for 10 min at 37°C, vortexed, then 12 min at 37°C. Separately, 2.2 mg of PTX was dissolved in 40 µL PEG 200, heated for 10 min at 80°C, vortexing at 2 min and 6 min. the fibrinogen solution was placed on ice water and allowed to cool (about 2 minutes). The PEG-PTX solution was removed from heat and allowed to cool to room temperature. 40 µL of PEG-PTX was added to the bottom of the fibrinogen solution tube (which is in ice water) with a manual pipette. The solution was sonicated in an ice water bath for 3 minutes at 4.5 power setting. 125 µL of 0.2M phosphate buffer pH 7.4 (no NaCl) was added to the solution and vortexed. The solution was sonicated for 3 minutes at 4.5 power (in an ice water bath). The solution was filtered through a 1 micron filter and measured by DLS. For injection, the appropriate volume was used depending on the weight of the mouse.

***Fibrinogen-Docetaxel Nanoparticle (FDN-DTX) Preparation:*** FDN-DTX nanoparticles were prepared using a sonication-based formulation method adapted for in vivo dosing and scaled to mouse body weight. For each mouse, a 2× volume was prepared to facilitate sonication of small volumes. Human fibrinogen was dissolved at a dose of 525 mg/kg (2×; e.g., 21 mg for a 20 g mouse) in a mixture of 33 µL 1 M Tris buffer (pH 8.0), 66 µL deionized water, and 200 µL PBS. The solution was incubated for ~10 minutes and vortexed to ensure complete dissolution. Docetaxel (DTX) was dissolved in PEG-200 at 50 mg/mL by heating to 80°C for 2 minutes, repeated three times, until fully solubilized. After approximately 20 minutes, once the fibrinogen solution was fully dissolved, the appropriate volume of DTX solution (corresponding to a final dose of 90 mg/kg, 2×) was added by pipetting to the bottom of the fibrinogen solution. The mixture was vortexed for 10 seconds and sonicated using a microtip probe sonicator at power level 3 for 3 minutes in an ice-water bath, taking care to avoid foaming. Following initial sonication, 400 µL PBS was added, and the solution was briefly vortexed and further sonicated at power level 4.5 for 4 minutes to ensure homogeneous nanoparticle formation. Nanoparticles were used within approximately 20 minutes of preparation and administered intravenously to deliver final doses of 525 mg/kg fibrinogen and 90 mg/kg docetaxel. Immediately after preparation, nanoparticles exhibited an average diameter of 433 ± 137 nm (polydispersity index [PDI] 0.22 ± 0.06). At 30 minutes post-preparation, particle size increased to 575 ± 124 nm with a PDI of 0.29 ± 0.05 (n = 6).

***EMT6 Triple-Negative Breast Tumor Model and Treatment:*** A syngeneic triple-negative breast cancer model was established using EMT6 murine mammary carcinoma cells. EMT6 cells ($1x10^6$ tumor cells mixed 1:1 with Matrigel in 30 µL final volume) were implanted subcutaneously into the hindlimbs of immunocompetent mice and allowed to grow until tumors reached the indicated starting volume prior to treatment. For treatment experiments, mice received FDN-PTX prepared as described above. Nanoparticles were administered intravenously at a fibrinogen dose of 525 mg/kg together with the indicated paclitaxel payload. In experiments employing vascular priming, mice received the vascular disrupting agent DMXAA (18 mg/kg, intraperitoneal) shortly before nanoparticle administration. Tumor growth was monitored by caliper measurements and the tumor volume was calculated using the formula: $V = 0.5(\text{length x width}^2)$. Animals were followed longitudinally for tumor regression, recurrence and survival.

***KPC Pancreatic Tumor Model and Treatment:*** A syngeneic pancreatic ductal adenocarcinoma model was established using KPC tumor cells derived from Kras^LSL-G12D/+; Trp53^LSL-R172H/+; Pdx1-Cre mice. Tumor cells ($2x10^6$) were mixed 1:1 with Matrigel in 100 µL final volume) and implanted subcutaneously into the hindlimbs of immunocompetent C57/BL mice and allowed to grow until tumors reached the indicated starting volume prior to treatment. For treatment experiments, mice received FDN-PTX or FDN-DTX. Nanoparticles were administered intravenously at a fibrinogen dose of 525 mg/kg together with the indicated taxane payload. Mice received the vascular disrupting agent DMXAA (18 mg/kg, intraperitoneal) 20 min before nanoparticle administration and 10 min after nanoparticle completion. Tumor growth was monitored by caliper measurements and the tumor volume was calculated using the formula: $V = 0.5(\text{length x width}^2)$. Animals were followed longitudinally for tumor regression, recurrence and survival.

***Clauss assay:*** Samples at the fibrinogen concentration indicated were mixed with 75 NIH U/mL thrombin and the absorbance at 405nm was measured over time.

***Histology:*** Tumors were excised 24 hours after treatment with VDA only (18 mg/kg, IP) or VDA (18 mg/kg, IP) plus FDNs (525 mg fibrinogen/kg, 44 mg PTX/kg, IV) and placed in formalin. After 48 hours the tumors were washed several times with PBS. Immunohistochemistry was performed by Premier Laboratory, LLC, Longmont, CO histology service and stained using anti-fibrinogen/fibrin antibodies (reactive with bovine and human) and secondary horse radish peroxidase and 3,3'-Diaminobenzidine (DAB). Positive and negative controls were done and VDA and VDA+ FDN tumors were processed on the same slide to avoid any differential staining.

***Drug delivery and pharmacokinetics:*** Tissue and blood samples, collected at specified times after IV injection of nanoparticles, were processed as published (*72*), with minor modifications. Briefly, frozen samples were mixed with and 10 volumes of 4% BSA in di$H_2O$ and placed in a ball mill (Retsch MM200) for 10 minutes at 30 Hz. 100 µL of homogenate was then removed and mixed with 10 mL ethyl acetate spiked with 100 µL of Butyl 4-hydroxybenzoate (2 ng/µL) in ethyl acetate as an internal standard (*73*). The solution was shaken vigorously and spun at 3000xg for 5 minutes. The organic layer was removed, transferred to a test tube in a 4°C water bath, and evaporated under a stream of air to complete dryness. The dried residue was reconstituted in 50 µL of acetonitrile, vortexed vigorously, and 50 µL DI water was added. The entire sample (100 µL) was loaded onto an Eclipse XDB-C18 column (5 µm, 3.0 x 150 mm, Agilent, 993967-302) and separated isocratically at 1 mL/min in 1:1 acetonitrile:water using a Agilent 1260 Infinity HPLC system. Peaks were integrated and normalized by the internal standard.  Glass materials instead of plastic were used throughout for HPLC experiments.

***Encapsulation Efficiency:*** 15 mg of fibrinogen was dissolved in 1 mL PBS. 1.5 mg of paclitaxel was dissolved in 40 uL PEG200 and heated at 80°C until fully dissolved. The fibrinogen was kept on ice for 15 minutes. 40 uL of PTX solution was added to the 1 mL PBS solution, vortexed for 5 seconds, returned to ice and sonicated on power 4.5 for 3 minutes. 500 uL of the fibrinogen-PTX solution (containing 0.75 mg PTX) was removed and 100 mg of ammonium sulfate was added. The suspension was vortexed for 2 minutes to completely dissolve the ammonium sulfate and precipitate the fibrinogen, and then spun at 16k x g for 2 minutes.  The supernatant was completely removed and 1 mL of ethanol was added to the pellet. The suspension was vortexed, sonicated briefly and spun at 16k x g for 2 minutes.  The supernatant was removed, diluted 1:50 in ethanol and the absorbance at 228 nm was recorded. The encapsulation efficiency was calculated as the ratio of OD 228 nm of the ethanol supernatant compared to OD 228 of 0.75 mg of PTX directly dissolved in ethanol and used as the value for 100% recovery.

***Drug release:*** A method previously described was used (*25*) with modifications. 5 µL of fibrinogen-PTX nanoparticle solution (containing 11 µg of PTX) was added to 95 µL of 10 mM phosphate buffer, pH 7.0. 10 units of thrombin were added, and the nanoparticle solution was allowed to clot at 37°C for 45 minutes. 1 mL of 4% BSA in 10 mM phosphate buffer was then gently added to the fully clotted solution, mixed completely with a Pasteur pipette, and placed back at 37°C to begin the release assay. 200 µL of supernatant were removed at 1, 15, 45 and 90 minutes after BSA addition, centrifuged 2 minutes at 16k x g, and extracted with 5 mL of ethyl acetate and processed as indicated in the drug delivery and pharmacokinetics section.

***Platelet aggregation microscopy study:*** Microscope slides were prepared by spreading a 0.1% poly-lysine solution and drying with a heat gun to enhance cell adhesion.  Blood was freshly drawn from a mouse and mixed with trisodium citrate anticoagulant (85 mM trisodium citrate, 71 mM citric acid, and 111 mM dextrose) in a ratio of 1 part anticoagulant to 9 parts blood. Platelet rich plasma (PRP) was obtained by centrifuging the blood at 200 x g for 10 minutes and taking the supernatant.  For assessment of activated platelets, 8 µL of PRP was applied to the slide and 8 µL of FDNs (26 mg/mL fibrinogen, 2 mg/mL PTX) and 1 µL of ADP (0.1mg/mL) were immediately added and stirred with a pipette tip..  The mixed drop was then spread over the slide and allowed to air dry. Staining was done by adding 6 drops of Wright-Giemsa stain 1 drop at a time with a 10 sec delay between drops. A drop of Permount was added and a coverslip applied. Slides were imaged at 1000x using oil immersion with a Nikon Optiphot-2 microscope with DS-F11 camera and NIS-Elements D3.2 software. For assessment of activation-inhibited platelets, 1 mL of blood was mixed with anticoagulant (as above) and 1 µL of a 1 mg/ml solution of prostaglandin E1. PRP was prepared as above then 8 µL PRP mixed with 8 µl of FDNs and the slide prepared as described above.

***Fluorescent fibrinogen nanoparticles:*** 14 mg bovine fibrinogen (Thermo Scientific J63276.06) was dissolved in 1 mL of 0.1 M phosphate buffer, pH 7.5. 0.4 mg of the lipophilic BODIPY 493/503 (Cas no. 121207-31-6, TCI Chemicals) was dissolved in 50 µL dimethyl sulfoxide (DMSO) and heated for 2 minutes at 80°C. When cooled to room temperature, the BODIPY was added to the fibrinogen solution and sonicated in an ice water bath using a microtip sonicator (Misonix Inc., model XL2000) with the tip in the solution at power setting 4.5 for 4 minutes. A clear transparent orange solution of nanoparticles resulted with a size of 314 nm and polydispersity of 0.28 as measured by dynamic light scattering using a Brookhaven Instruments Corp. 90Plus machine. Some of the solution was diluted 50-fold and exposed to a 365 nm UV lamp (Spectroline model ENF-260C) and photographed.

***Institutional animal assurance:*** Animal experiments were conducted according to NIH guidelines and approved by Nanoprobes' Institutional Animal Care and Use Committee before start of the study.

### Supplementary Text

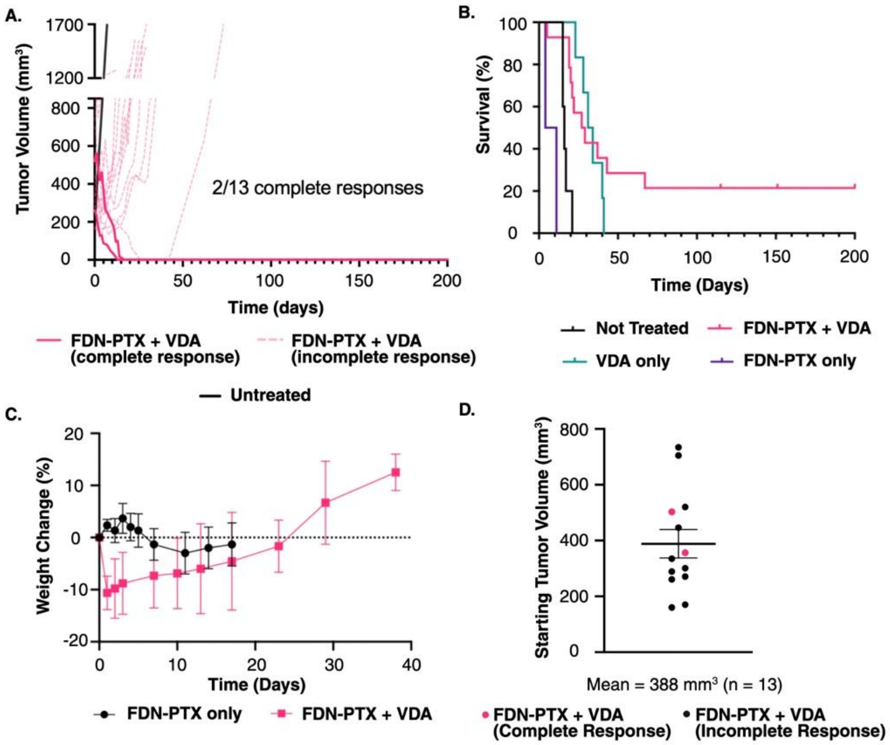


**Fig. S1 Paclitaxel-loaded fibrinogen nanoparticles produce heterogeneous responses in KPC pancreatic tumors.** **(A)** Individual tumor growth trajectories following treatment with fibrinogen–paclitaxel nanoparticles (FDN-PTX) combined with a vascular disrupting agent (VDA) in KPC tumor–bearing mice (n = 13). Two of thirteen treated animals exhibited complete tumor regression (solid pink lines), whereas the remaining animals showed partial or transient responses followed by tumor progression (dashed pink lines). Representative untreated tumor growth is shown for comparison. **(B)** Kaplan–Meier survival analysis corresponding to panel A. VDA + FDN-PTX improved survival relative to untreated controls (n = 5) but did not produce consistent durable tumor eradication. **(C)** Body weight changes following treatment with FDN-PTX alone (n = 3) or FDN-PTX + VDA (n = 13). Combination therapy produced a transient decrease in body weight during the first week followed by recovery to baseline. **(D)** Distribution of starting tumor volumes in the FDN-PTX + VDA cohort (n = 13). Tumors ranged broadly in size at treatment initiation (mean = 388 mm³). Pink points denote mice that achieved complete tumor regression.
